# Complex epidemiological dynamics driven by the combination of host spatial structure and seasonal forcing

**DOI:** 10.64898/2026.08.10.743859

**Authors:** Alex Best, Andy White, Mike Boots

**Affiliations:** School of Mathematical and Physical Sciences, University of Sheffield, Sheffield, UK; Department of Mathematics and Computer Science, Heriot-Watt University, Edinburgh, UK; Integrative Biology, University of California Berkeley, Berkeley, CA, USA

## Abstract

Spatial population structure and seasonality are both central to the spread of many infectious diseases of plants, animals and humans. While seasonal forcing in transmission often plays an important role in epidemiological models of a wide range of infectious disease, and we now have some theoretical understanding of the dynamical impacts of spatial structure, the combined effects of these two ubiquitous processes has not been examined in detail. Here, we develop a novel model to explore the combined influence of spatial structure and temporal variability on disease dynamics. Spatial structure is represented using a lattice-based approach with near-neighbour interactions, while temporal variability is included through regular, seasonal, variation of the transmission rate. We use bifurcation analysis of a pair approximation of the full spatial model to identify the parameter regimes associated with qualitatively distinct dynamical behaviours. The model exhibits a remarkably wide range of complex dynamics, including limit cycles, quasi-periodic cycles, multi-year cycles, chaotic dynamics and bistability between these different states. In particular, complex dynamics occur when reproduction is predominantly local, with the dynamics depending critically on the amplitude of the seasonal transmission rate. We show how high transmission rates, high birth rates and in particular low recovery rates are requirements for complex dynamics. We predict that ‘SI’-type disease interactions in plant pathogen systems will show complex dynamics even with relatively global transmission dynamics.

## 1 Introduction

Epidemiological dynamics are fundamentally spatio-temporal in nature. Individual hosts must come into contact with one another, directly or indirectly, for successful transmission of disease, meaning where hosts are located at any particular time is crucial [Lion and van Baalen, 2008]. Additionally, the life histories of both hosts and the parasites that infect them can be highly dependent on the climate and wider environment, resulting in substantial temporal variation in population size and life-history traits [Altizer et al., 2006]. Clearly, diseases of plants are subject to particularly strong spatial and temporal variation as they are more directly shaped by their natural environment, with a range of empirical and statistical studies highlighting the spatiotemporal dynamics of plant diseases [Reynolds et al., 1988, Madden and Hughes, 1995, Gottwald et al., 2002]. Furthermore, Cunniffe et al. [2015] list the inclusion of host spatial structure and temporal variation as two of the key challenges for the plant disease modelling field. Spatiotemporal features are not limited to plant diseases, however, but are also central to diseases of animals and even humans; for example, a number of recent studies have highlighted the spatiotemporal nature of the spread of COVID-19 [Castro et al., 2021, Kim et al., 2021, Jalal et al., 2024]. Understanding the combined effects of spatial structure and temporal variation is therefore a central question for epidemiological models.

A variety of techniques exist for including spatial structure in disease models, including metapopulation frameworks with migration between patches [e.g. Wadkin et al., 2023], dispersal kernel approaches [e.g. Cunniffe et al., 2014] and network models [Keeling and Eames, 2005] Our focus here is on a particular version of a network model, namely a lattice-based framework with a pair approximation [Matsuda et al., 1992, Keeling, 1999]. In this approach, host individuals are assumed to exist on a regular lattice. Processes such as infection and host birth can be assumed to be either ‘global’ (mean-field) or ‘local’ (near-neighbour) or some combination of the two. This system can be expressed as a set of ordinary differential equations that follow not only the proportions of ‘singlet’ densities (i.e. susceptibles or infecteds) but also ‘pair’ densities (i.e. susceptible-infected neighbours). The initial system is not closed, but a pair approximation can then be applied to produce a closed model [Matsuda et al., 1992, Sato et al., 1994]. This approach has been used by a number of studies to explore the epidemiological and evolutionary impact of spatial structure [Boots and Sasaki, 1999, Filipe et al., 2004, Kamo and Boots, 2006, Webb et al., 2007a,b, Lion and Boots, 2010, Best et al., 2011, 2012, Débarre et al., 2012, Maltz and Fabricius, 2016, Wren and Best, 2021, Best and Cunniffe, 2026]. For example, Wren and Best [2021] showed in a basic SIR model that increasing the proportion of local infections has a non-linear impact on the overall disease dynamics, with high degrees of local infection leading to lower and later epidemics. From a more ecological perspective, where host demography is included, Webb et al. [2007a] and Webb et al. [2007b] showed that local infection and reproduction can lead to endemic limit cycles and disease-driven extinction - outcomes that do not occur in the standard mean-field SIR model - when the transmission rate and birth rate are high. Recent work by Best and Cunniffe [2026] further showed that such a lattice-based model can provide an excellent fit to real-world experimental data for the spread of an economically important plant disease, predicting high levels of localised spread. Moreover, studies using dispersal kernel approaches have also demonstrated good fits to experimental data of plant pathogens [Meentemeyer et al., 2011, Neri et al., 2014, Cunniffe et al., 2014], highlighting the importance of including spatial structure and localised disease spread to model these real-world systems.

There is a rich history of epidemiological models exploring the impacts of seasonality on disease spread, particularly in childhood infectious diseases, where transmission is strongly influenced by school terms. [Fine and Clarkson, 1982, Earn et al., 2000]. Seasonal forcing has also been extensively studied in ecological host–parasite systems, where it is recognised as a major driver of disease dynamics[White et al., 1996, Ireland et al., 2004, Smith et al., 2008, Best, 2013, Taylor et al., 2015, Ward and Best, 2021]. A well-known result from seasonal models is that increasing the amplitude of periodic forcing can lead to a period-doubling cascade to chaos [Aron and Schartz, 1984, Schwartz, 1985]. Furthermore, studies often find the existence of multiple stable cycles, potentially of different periods, for the same parameter sets [Earn et al., 2000, Taylor et al., 2015].

Relatively few mechanistic modelling studies have incorporated both spatial structure and seasonality. Those that do are often individual-based computational models tailored to specific pathosystems. In plants, examples include potato late-blight [Skelsey et al., 2010], sudden oak death [Meentemeyer et al., 2011] and olive quick-decline syndrome [Chapman et al., 2025], in animals, foot and mouth disease in deer [Highfield et al., 2009], raccoon rabies [Duke-Sylvester et al., 2010], and chronic wasting disease in cervids [Oraby et al., 2014], and in humans, influenza-A [Abed et al., 2026], and COVID-19 [Thornton et al., 2026]. While undoubtedly useful for predicting the epidemiological dynamics in specific circumstances, these models provide less general biological insight into the joint roles of spatial structure and seasonality. From a more theoretical perspective, a small number of studies have examined the mathematical properties of models with infection spread through a reaction-diffusion process and seasonal transmission [Peng and Zhao, 2012, Liu et al., 2019], largely focused on defining disease persistence properties such as *R*_0_. Consequently, we have little general understanding of how spatial structure and seasonality - two central components of most real-world disease systems - jointly interact to shape epidemiological dynamics.

Our aim is to investigate the dynamics of an epidemiological model that includes both spatial structure and seasonality. Specifically, we take a lattice-based approach that novelly represents infection spread as a mix of near-neighbour and mean-field transmission, and that includes sinusoidal variation in the transmission rate. To characterise the resulting dynamics, we adopt a bifurcation analysis approach, using the numerical continuation software AUTO-07p [Doedel and Oldeman, 2009]. Given the ubiquity of both spatial structure and seasonal forcing in many important disease systems, our results are important to better understand the range of dynamics that may arise in real-world disease systems.

## 2 Methods

Our model follows a Susceptible - Infected - Susceptible (SIS) compartmental framework with births and deaths. Let *P*_*S*_ and *P*_*I*_ be the probabilities that, if we choose a random site on the lattice, it is occupied by a susceptible or infected individual, respectively. As we include deaths there are also potentially empty sites on the lattice, *P*_0_, where *P*_*S*_ + *P*_*I*_ + *P*_0_ = 1. As such, we need only track two of these variables with differential equations. We also define nearest neighbour terms, such as *q*_*I/S*_, to be the conditional probability that if we choose a susceptible individual, it has a neighbour that is infected, and similarly for other combinations. We can describe the dynamics of the ‘singlet’ densities by the following equations,

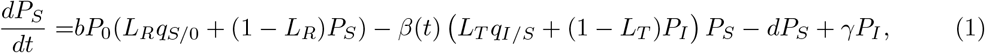

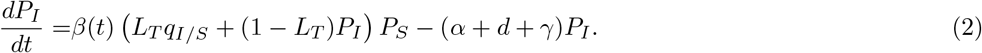

The birth rate is *b*, the host mortality is *d*, the parasite-induced virulence is *α* and host recovery is *γ*. Only susceptible hosts reproduce and all offpring are born susceptible. The parameter *β*(*t*) gives the rate of infection from infected individuals. This is assumed to fluctuate sinusoidally over the course of a year, with the functional form *β*(*t*) = *β*_0_[1+ *δ* sin(2*πt*)], where *β*_0_ is the average transmission and *δ* is the amplitude of the oscillations. The parameters *L*_*T*_, *L*_*R*_ ∈ [0, 1] control the respective ratios of local-to-global transmission and reproduction. We assume ‘local’ infections/births are purely near-neighbour, and ‘global’ infections/births are random (mean-field).

This system is not closed, since, for example, *q*_*I/S*_ = *P*_*SI*_*/P*_*S*_, where *P*_*SI*_ is the probability that if we choose a neighbouring pair of sites, one is susceptible and the other infected. We therefore need to write down equations describing the dynamics of these ‘pair’ densities. We can find these by considering the possible transitions between pairs of sites. For example, consider the creation and loss of an [*S, I*]-pair:

### created

- [*S, S*] → [*S, I*] one member of an [*S, S*]-pair becomes infected from:
  - another *I* neighbour (since each individual has 4 neighbours) at rate 0.75*βL*_*T*_ *q*_*I/SS*_
  - a random *I* contact at rate *β*(1 − *L*_*T*_)*P*_*I*_
- [0, *I*] → [*S, I*] a newborn susceptible appears in the empty site of an [0, *I*]-pair, from:
  - another S neighbour at rate 0.75*bL*_*R*_*q*_*S/*0*I*_
  - a random *S* contact at rate *b*(1 − *L*_*R*_)*P*_*S*_
- [*I, I*] → [*S, I*] one member of an [*I, I*]-pair recovers at rate *γ*

### lost

- [*S, I*] → [*I, I*] *S* member of the pair becomes infected from
  - its known *I* neighbour (i.e. from the [*S, I*]-pair) at rate 0.25*βL*_*T*_
  - another *I* neighbour (since each individual has 4 neighbours) at rate 0.75*βL*_*T*_ *q*_*I/SI*_
  - a random *I* contact at rate *β*(1 − *L*_*T*_)*P*_*I*_
- [*S, I*] → [*S, S*] *I* member of the pair recovers at rate *γ*
- [*S, I*] → [0, *I*] *S* member of the pair dies at rate *d*
- [*S, I*] → [*S*, 0] *I* member of the pair dies either from background mortality at rate *d* or parasite-induced virulence at rate *α*.

Considering all such possible transitions we reach the following set of equations,

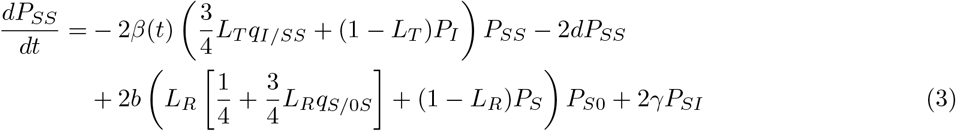

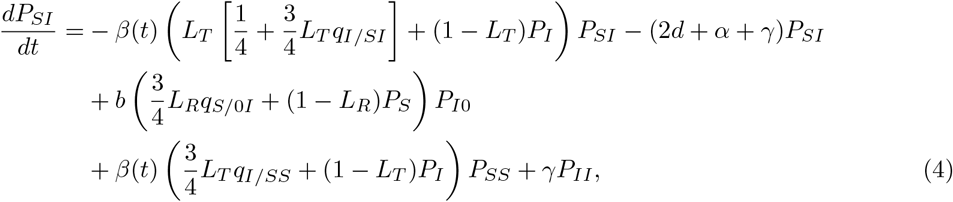

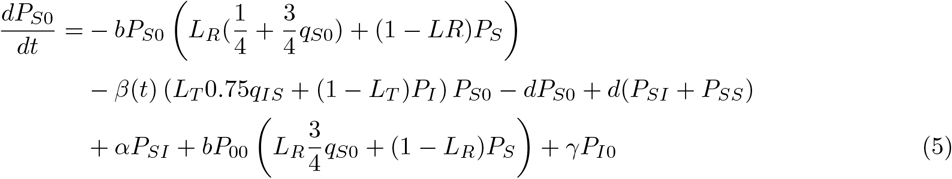

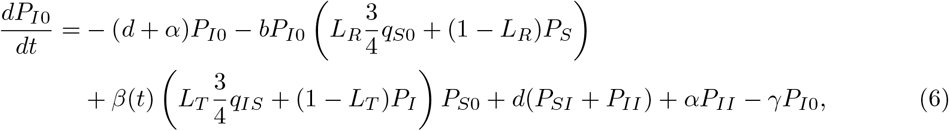

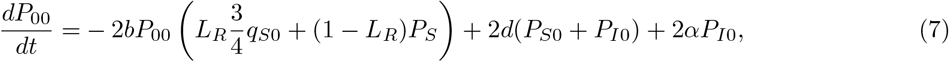

with *P*_*II*_ = 1 − (*P*_*SS*_ + *P*_00_) − 2(*P*_*SI*_ + *P*_*S*0_ + *P*_*I*0_). This system is also not closed, since we have conditional probabilities such as *q*_*I/SS*_ = *P*_*SSI*_*/P*_*SS*_, which would require writing down the dynamics of ‘triplet’ densities. At this point we can apply a pair approximation, letting *q*_*I/SS*_ = *q*_*I/S*_, which closes the system [Matsuda et al., 1992].

Our primary method for analysis of the model is to take a bifurcation approach using the numerical continuation software AUTO-07p [Doedel and Oldeman, 2009]. This allows us to identify bifurcations in the model as one or more parameters are varied, and therefore divide parameter space into regions of qualitatively different epidemiological dynamics. We found that the system was often stiff, meaning both AUTO-07p and numerical simulations in Python failed to run. We therefore log-transformed the above model before running, transforming back to the true variables before plotting the outputs. We will expect to see a number of different types of bifurcation in our model which we briefly detail here:

- **Transcritical bifurcation** - where two equilibria collide and swap stability. In epidemiological models the most common example is where the disease-free equilibrium loses stability and the stable endemic equilibrium emerges.
- **Period-doubling bifurcation** - where the period of limit cycles doubles, usually as an integer multiple of the forcing period.
- **Fold bifurcation** - often called a saddle-node or ‘blue sky’ bifurcation, this occurs when two steady states - usually a saddle and a node - appear together and diverge, causing a fold in the solution branch. This commonly causes a region of bistability, with the saddle separating the stable node and some other steady state.
- **Torus bifurcation** - this gives rise to quasi-periodic cycles, whose trajectories wrap around a torus, never quite completing a closed orbit. They usually map on to a Hopf bifurcation when there is no periodic forcing.

We have provided code on Github that readers may use to reproduce our results, at https://github.com/abestshef/seaso

## 3 Results

### 3.1 Fully local model

We begin by setting both *L*_*T*_ = 1 and *L*_*R*_ = 1 such that the model is fully local, and we also initially set *γ* = 0 meaning this is an SI model. We choose a default parameter set with relatively high birth and transmission rates (*b* = 60, *β* = 60) and explore the impact of introducing and increasing the amplitude of seasonal forcing in a bifurcation diagram (figure 1). In the absence of seasonality, for our chosen parameter set, there are limit cycles. As forcing is introduced these morph into quasi-periodic, or toroidal, cycles - non-periodic and non-repeating yet somewhat predictable oscillations with a dominant non-integer period. As the amplitude is increased, the system passes through a torus bifurcation where the system briefly stabilises into regular 1-year cycles. There are then two fold bifurcations in the solution branch, meaning most of this low amplitude region is in fact bistable: alongside the quasi-periodic cycles and 1-year cycles just mentioned, there are additional stable 1-year cycles. As the amplitude is increased further there is eventually a period-doubling bifurcation and 2-year cycles emerge. A further period-doubling to 4-year cycles then occurs and remains stable up to the maximum amplitude. We show sample dynamics from each of these regions in figure 2, including demonstrating the bistability where two cases have the same parameter values but different initial conditions. In particular we see non-forced limit cycles (*δ* = 0), quasi-periodic cycles (*δ* = 0.015), 1-year cycles (*δ* = 0.015 and *δ* = 0.5), 2-year cycles (*δ* = 0.8) and 4-year cycles (*δ* = 0.95).

**Figure 1:**
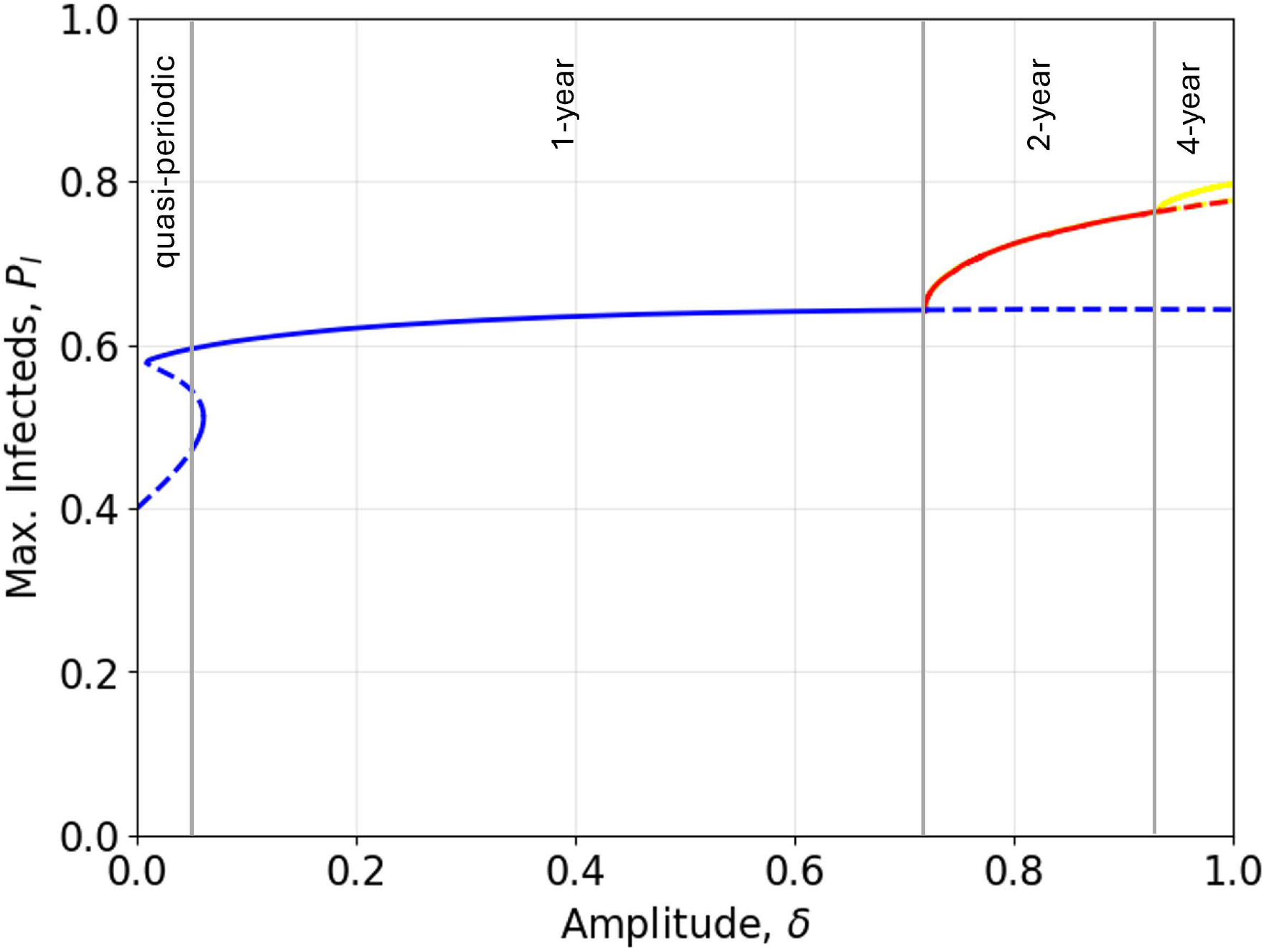
A bifurcation diagram as the amplitude, *δ* is varied. Solid lines denote stable branches and dashed lines are ustable branches. The blue curve is the principal (1-year cycle) solution, the red curve the 2-year branch and the yellow curve the 4-year branch. The grey vertical lines partition the regions of different outcomes as labelled. *b* = 60, *d* = 1, *β* = 60, *α* = 1, *γ* = 0, *L*_*T*_ = 1, *L*_*R*_ = 1.

**Figure 2:**
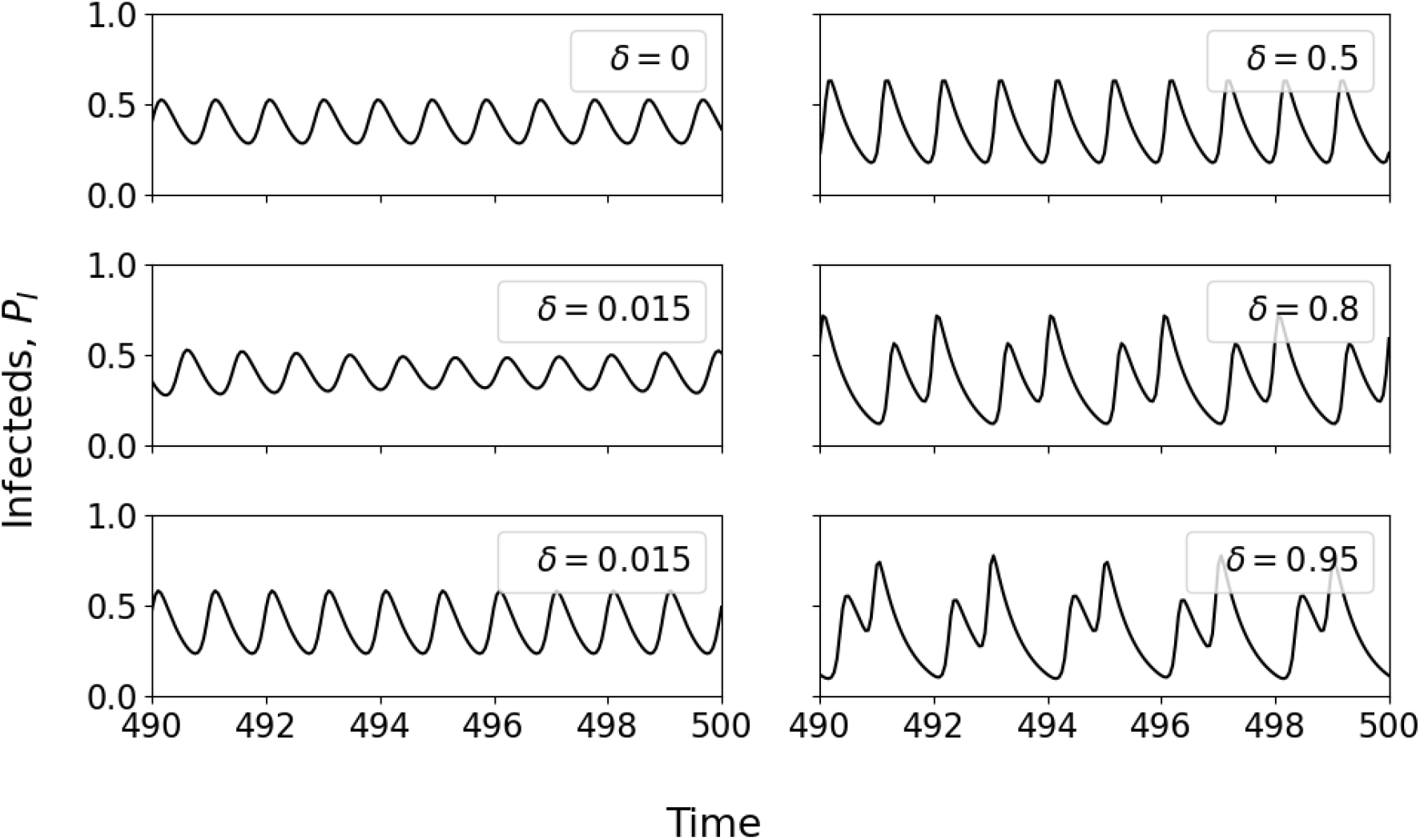
A selection of numerical simulations of the model for different values of amplitude, *δ*, chosen to demonstrate the different outcomes found in figure 1. *b* = 60, *d* = 1, *β* = 60, *α* = 1, *γ* = 0, *L*_*T*_ = 1, *L*_*R*_ = 1.

We investigate the system further with a co-dimension 2 bifurcation diagram, tracking the bifurcation curves when both the amplitude and the transmission rate are varied. Figure 3 shows that (once transmission is high enough for the parasite to persist) one-year cycles remain the norm up until moderately high rates of infection. However, there is then a rich array of possible dynamics at higher transmission rates. If the forcing amplitude is low, there is bistability, either between quasi-periodic and 1-year cycles or two different 1-year cycles. When the amplitude is high then we see not only the 2-year and 4-year cycles found in figure 1, but a full period-doubling cascade towards chaotic dynamics.

**Figure 3:**
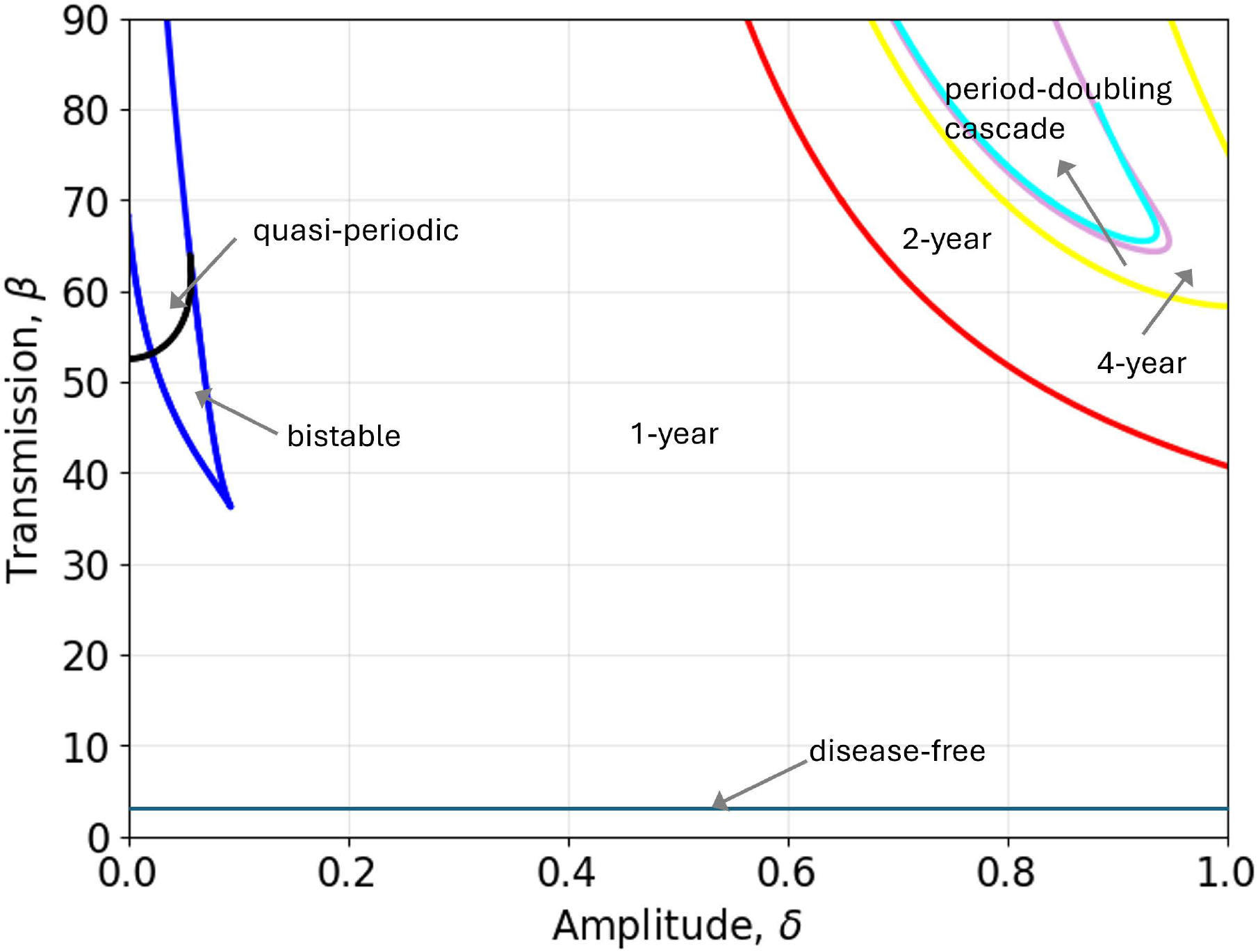
Co-dimension 2 bifurcation diagram of the outcomes as transmission, *β* and amplitude *δ* are varied. The thin black line marks a transcritical bifurcation, the thick black line is a torus bifurcation, the blue line is a fold bifurcation and all other lines are period-doubling bifurcations. *b* = 60, *d* = 1, *α* = 1, *γ* = 0, *L*_*T*_ = 1, *L*_*R*_ = 1.

### 3.2 Effects of local interactions

We now take a detailed look at the impact of varying the degrees of local transmission and local reproduction by making *L*_*T*_ and *L*_*R*_ our bifurcation parameters. We initially study how each interacts with the amplitude of the forcing, setting the other parameter to be fully local. Figure 4a shows that unless reproduction is predominantly local, the system stabilises to regular 1-year cycles. If *L*_*R*_ is high, we see similar behaviour as in figure 3: for low amplitude forcing there are quasi-periodic cycles and bistability, while for high amplitude forcing there are 2-year cycles.

**Figure 4:**
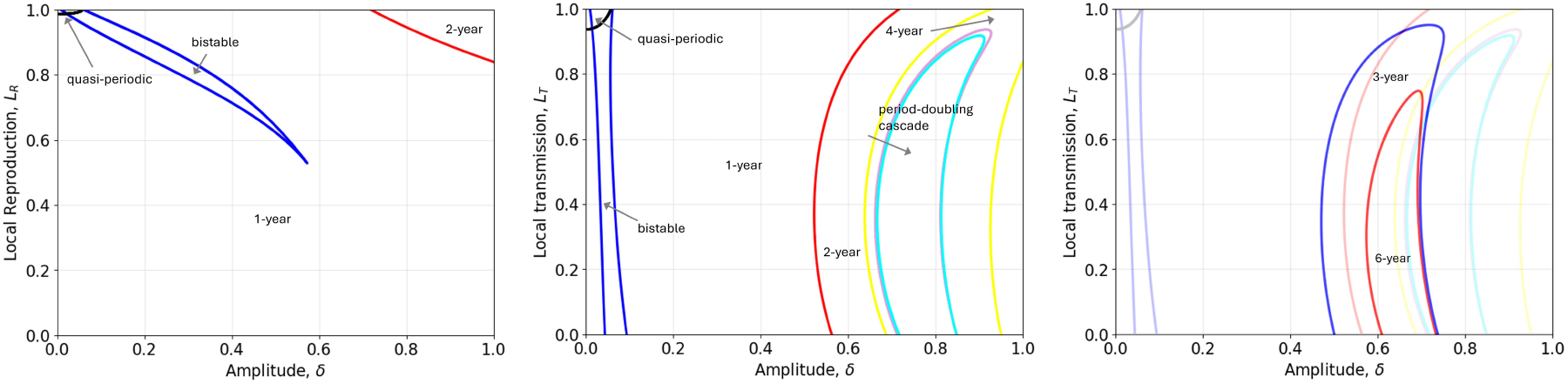
Bifurcation diagrams as the local parameters are co-varied with the amplitude. Each curve marks a period-doubling bifurcation. (a) Local reproduction, *L*_*R*_ and amplitude, *δ*. (b) and (c) Local transmission, *L*_*T*_, and amplitude, *δ*. Default parameter values: *β* = 60, *b* = 60, *L*_*R*_ = 1, *L*_*T*_ = 1, *d* = 1, *α* = 1, *γ* = 0.

The behaviour as local transmission, *L*_*T*_ is varied with the amplitude of forcing is rather more complex (figure 4b). Here, the multi-period cycles can occur for even fully global transmission (we note that *L*_*R*_ = 1 here). As before, low amplitude cycles tend to produce bistability (and when *L*_*T*_ is high, quasi-periodic cycles). Now high amplitude forcing produces not only 2-year cycles but a full period-doubling cascade towards chaotic dynamics. While conducting confirmatory simulations of the system, we noticed the presence of an additional 3-year cycle branch that is not on the main solution branch. We ran further AUTO continuations from this cycle, revealing that for intermediate and low *L*_*T*_ and intermediate *δ*, 3-year and 6-year cycles are also possible, as shown in figure 4c (overlain with the original plot from 4b).

Finally we examine the behaviour as the two local terms co-vary. We deliberately choose default parameters from figure 3 that place us close to the period-doubling bifurcations, with *β* = 60 and *δ* = 0.75. For ease, we do not track the 3-year branch identified in figure 4c but note it would be present here. Figure 5a reinforces that local reproduction, *L*_*R*_, has an especially strong impact on the dynamics, with regular 1-year cycles always found below a threshold value of *L*_*R*_. However, as the local proportion of reproduction is increased there is a period-doubling cascade resulting in high-period, near-chaotic cycles when *L*_*R*_ is high. In contrast, increasing local transmission *L*_*T*_ slightly stabilises the dynamics, with the most complex cycles requiring high *L*_*R*_ but intermediate or low *L*_*T*_ . We demonstrate a number of the different types of cycles from simulations of the ODEs in figure 5b, namely 1-year (*δ* = 0.6), 2-year (*δ* = 0.85), 4-year (*δ* = 0.95) and 16-year (*δ* = 0.97) cycles.

**Figure 5:**
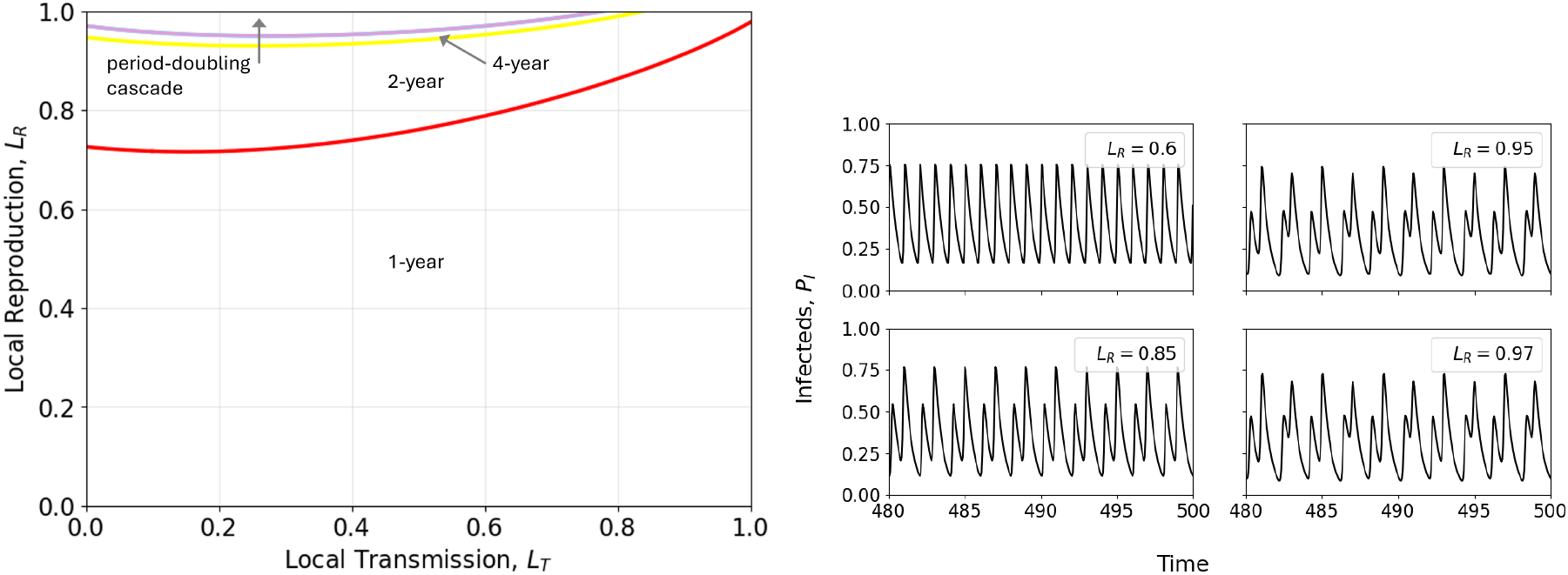
(a) Bifurcation diagrams as the two local parameters are varied. Each curve marks a period-doubling bifurcation. At high values of *L*_*R*_ the 4-yr//8-yr and 8-yr/16-yr almost overlay one another as a period-doubling cascade rapidly occurs. (b) Simulations of some of the dynamics in panel (a), with *L*_*T*_ = 0.5 and the values of *L*_*R*_ as shown in the legends. Default parameter values: *β* = 60, *δ* = 0.75, *b* = 60, *d* = 1, *α* = 1, *γ* = 0.

The results we have presented so far have used the pair approximation model of equations (1)-(7), which, as discussed in the methods, does not fully incorporate the finer spatial details as it removes higher-order correlations from the equations. To confirm our findings, in particular on the impacts of the local parameters *L*_*R*_ and *L*_*T*_, we also developed fully spatially explicit simulations of the system using the well-known direct method stochastic simulation algorithm of Gillespie [1977]. This is an individual-based stochastic model where the event rates according to the equations in (1)-(7) are used to determine the time to the next event, drawn from an exponential distribution. Which event occurs then depends probabilistically on the relative value of each event rate. We use a 200 x 200 grid and employ tau-leaping when population sizes are large to speed up the simulations [Gillespie, 2001]. Figure 6 shows the resulting stochastic dynamics (red) alongside those of the pair approximation model (grey) for a selection of *L*_*R*_ and *L*_*T*_ values from figure 5a. These match up well qualitatively, in particular showing the predicted regular 1-year cycles with low *L*_*R*_, but more variable dynamics when *L*_*R*_ is high and 2-year or 4-year cycles are predicted. We note that in some runs, especially for large amplitude 1-year cycles and relatively small grid sizes, the susceptible population is prone to go extinct as its density falls close to 0 for prolonged periods.

**Figure 6:**
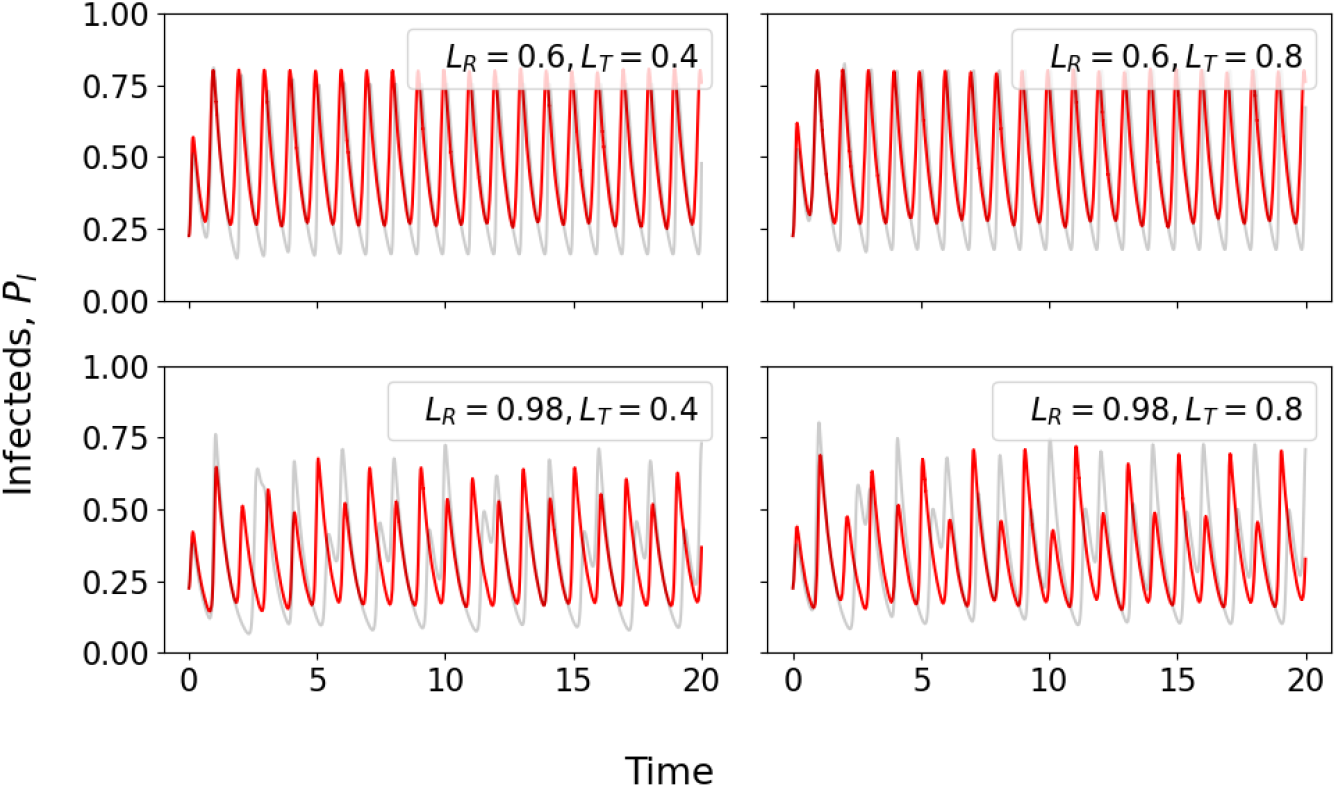
Output from stochastic simulations (red) overlaid with the predictions from the pairs model (grey). Default parameter values: *β* = 60, *δ* = 0.75, *b* = 60, *d* = 1, *α* = 1, *γ* = 0.

### 3.3 Effects of ecological & epidemiological parameters

Next we investigate how the ecological and epidemiological parameters impact the dynamics, again choosing our default values of *β* = 60 and *δ* = 0.75. Figure 7a shows a co-dimension 2 bifurcation diagram as the birth and mortality rates of the host are varied. When both are high (short-lived, fast reproducing hosts), the dynamics are regular 1-year cycles. However, for low death rates and intermediate to high birth rates (long-lived, fast reproducing hosts) then 2-year cycles emerge, with a further branch to 4-year cycles at the extreme end of low death rates and high birth rates. We note that the period of our seasonality is always one year. As such, this result suggests that multi-annual cycles might be more likely in hosts that live for multiple years.

**Figure 7:**
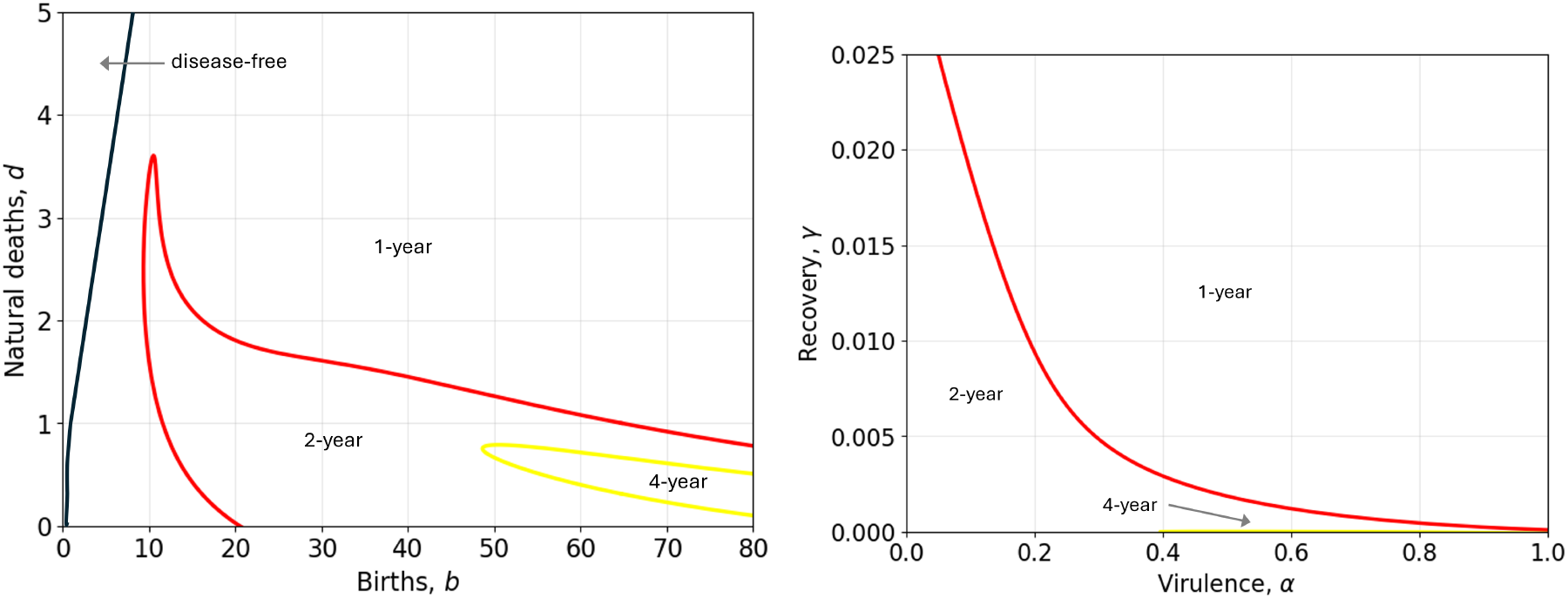
Bifurcation diagrams as model parameters are varied. (a) Co-varying births, *b* and deaths, *d*, (b) co-varying virulence *α* and recovery, *γ. β* = 60, *δ* = 0.7, *b* = 60, *d* = 1, *α* = 1, *γ* = 0, *L*_*T*_ = 1, *L*_*R*_ = 1.

Finally we co-vary two epidemiological traits, parasite virulence (*α*) and recovery (*γ*) in figure 7b. Relatively low rates of recovery cause the stabilisation of the system to 1-year cycles, with near-zero recovery required for cycles of 4-years or longer. This suggests complex dynamics are more likely in obligate killing parasites or hosts with limited immune defences. Multi-year cycles are also more likely for lower virulence rates.

## 4 Discussion

We have investigated the interplay of two important features of real-world disease systems - spatial structure and seasonality - each of which is known to cause complex epidemiological dynamics, but whose combination has not been comprehensively investigated. We have found that a remarkably rich array of epidemiological dynamics can arise when both features are included, including multi-annual cycles, quasi-periodic cycles and multi-stability. Moreover, we show that the requirements for the more complex dynamics are: highly local reproduction, moderately local transmission, high transmission rates, high birth rates, low death rates, low virulence and very low recovery. Furthermore, the amplitude of seasonal forcing plays a critical role in shaping the nature of this complexity: low amplitudes lead to quasi-periodic cycles and bistability, high amplitudes to multi-year cycles. In particular, our work highlights that infectious diseases in highly spatially structured hosts - such as many plant systems - where there is little recovery may show highly complex dynamics even if transmission is relatively global. There is likely to be much less effect in infectious diseases of relatively well mixed hosts - such as humans - with high rates of recovery.

Our results are particularly important to plant systems, which often have predominantly local dispersal/reproduction [Bullock et al., 2017] and seasonal life histories [Chunie and Régnière, 2017]. Indeed, Cunniffe et al. [2015] identify the need to understand the interplay between host spatial structure and temporal variation as a major challenge in plant disease modelling. Plant pathogens predominantly exhibit local transmission with occasional long-distance dispersal, often mediated by wind or rain. For example, infections of *Phytophthora ramorum*, the causal agent of sudden oak death, typically occur within 300m of an infected tree, although transmission over distances of up to 4 km has been documented [Hansen et al., 2008]. Similarly, asexual conidia of *Dothistroma septosporum*, the cause of needle blight in pine trees, generally disperse less than 200 yards [Gibson et al., 1964], but dispersal beyond 2 km has been observed [Mullett et al., 2016], while sexually produced ascospores are thought to travel much farther. A similar dichotomy of dispersal distances of a few metres for conidia and many kilometres for ascospores has been found in *Cryphonectria parasitica*, the cause of chestnut blight [Rigling and Prospero, 2018]. Mechanistic models fitted to epidemic data have shown that transmission is dominated by short-range dispersal [Meentemeyer et al., 2011, Neri et al., 2014, Cunniffe et al., 2014, Chapman et al., 2025]. For example, a recent study that used a lattice structure to model Bahia bark scaling of citrus estimated the local transmission parameter, *L*_*T*_, to lie between 0.93 and 1.0 [Best and Cunniffe, 2026], while studies using separate short-range and long-range dispersal kernels have estimated that over 90% of transmission occurs locally [Meentemeyer et al., 2011]. These findings suggest that many plant diseases satisfy the condition of intermediate local transmission that our study identifies as a driver of complex dynamics. Evidence for multi-year outbreaks is more limited because long-term datasets are rare, but records of annual potato late blight epidemics reveal both annual and multi-year cycles [Zwankhuizen and Zadoks, 2002]. Our results provide a mechanistic explanation for how variation in host life history and pathogen transmission could generate such diverse epidemic dynamics.

The combination of spatial structure and seasonality is by no means limited to plant diseases. Strong spatiotemporal signals have also been reported in diseases of animal [Osnas et al., 2009, Barasona et al., 2014] and human hosts [Chen et al., 2018, Castro et al., 2021]. While the nature of spatial structure is likely to differ from the assumptions made here, we suggest that the broad conclusions should still hold: more complex epidemiological dynamics arise in systems characterised by high transmission rates, predominantly localised birth and transmission, and long infectious periods. However, we find that the recovery rate is the most restrictive for the emergence of complex dynamics, with even relatively low recovery rates causing the system to stabilise to 1-year cycles. Both Webb et al. [2007a] and Webb et al. [2007b] similarly showed that in the non-seasonal but spatially-structured model, recovery tended to remove limit cycles. The same pattern was found by Ward and Best [2021] for a mean-field but seasonal epidemic model - higher recovery tended to stabilise the dynamics towards 1-year or 2-year cycles from more complex and sometimes chaotic dynamics. The influx of susceptible hosts due to recovery stabilises the system, preventing both the peak and trough of infected numbers from being too extreme, and therefore preventing multi-year cycles from emerging. Webb et al. [2007a] also show that more localised reproduction and transmission make limit cycles more likely in the non-seasonal model. Taken as a whole this suggests that our results are most likely to apply to human and animal diseases where there is little recovery. It’s an open question whether the spatial structure in these ‘SI’ type disease interactions is sufficient to lead to complex epidemiological dynamics when combined with seasonal forcing.

To incorporate temporal variation we have used a simple sinusoidal function, in common with many other modelling studies [Aron and Schartz, 1984, Schwartz, 1985, Ireland et al., 2004, Ward and Best, 2021]. Such a function approximates yearly changes in climate variables such as temperature or rainfall, especially in temperate zones. Of course, more realistic, data-derived functional forms could be used. For example, it is known that many plant diseases’ life-cycle depends on the ‘thermal time’ - the sum of degrees above a threshold over time [Lovell et al., 2004, Madden et al., 2007, Carisse et al., 2008, Zearfoss et al., 2011], and some mathematical and simulation models of specific pathosystems have used such a function for their time-varying transmission term [Kendall et al., 1992, Van Den Berg et al., 2016, Salotti and Rossi, 2023]. However, while such refinements may improve predictive accuracy in particular systems, they add complexity that is not appropriate for the more generic setting considered here. Models with seasonality but not spatial structure commonly show that as the amplitude of seasonality increases, more complexity arises [Aron and Schartz, 1984, Best, 2013]. These previous works lend weight to our findings for when complex epidemiological cycles are most likely when spatial structure and seasonality are combined.

In summary, our results suggest that the complex epidemiological dynamics that are seen in many plant disease systems may be due to the combination of seasonal forcing and strong host spatial structure. It’s important to note that there is often relatively little recovery in these systems and that our results suggest this is critical to the generation of rich epidemiological dynamics. In principle, ‘SI’ type infections in animal and human populations may also lead to more complex dynamics if there is sufficient spatial structure in the population, but the high recovery rates of many important infectious disease tends to reduce these effects.

## Acknowledgements

Many thanks to Jonathan Sherratt for helpful discussions on using AUTO-07p and to Jack Woodruff for feedback on the model. AB was funded in this work by an EPSRC Mathematical Sciences Small Grant UKRI1110. A.W. was supported by the BBSRC EEID research grant BB/V00378X/1. M.B. was supported by NSF-DEB-2011109 EEID research grant.

